# Experimental evolution reveals favored adaptive routes to cell aggregation in yeast

**DOI:** 10.1101/091876

**Authors:** Elyse A. Hope, Clara J. Amorosi, Aaron W. Miller, Kolena Dang, Caiti Smukowski Heil, Maitreya J. Dunham

## Abstract

Yeast flocculation is a community-building cell aggregation trait that is an important mechanism of stress resistance and a useful phenotype for brewers; however, it is also a nuisance in many industrial processes, in clinical settings, and in the laboratory. Chemostat-based evolution experiments are impaired by inadvertent selection for aggregation, which we observe in 35% of populations. These populations provide a testing ground for understanding the breadth of genetic mechanisms *Saccharomyces cerevisiae* uses to flocculate, and which of those mechanisms provide the biggest adaptive advantages. In this study, we employed experimental evolution as a tool to ask whether one or many routes to flocculation are favored, and to engineer a strain with reduced flocculation potential. Using a combination of whole genome sequencing and bulk segregant analysis, we identified causal mutations in 23 independent clones that had evolved cell aggregation during hundreds of generations of chemostat growth. In 12 of those clones we identified a transposable element insertion in the promoter region of known flocculation gene *FLO1*, and in an additional five clones we recovered loss-of-function mutations in transcriptional repressor *TUP1*, which regulates *FLO1* and other related genes. Other causal mutations were found in genes that have not been previously connected to flocculation. Evolving a *flo*1 deletion strain revealed that this single deletion reduces flocculation occurrences to 3%, and demonstrated the efficacy of using experimental evolution as a tool to identify and eliminate the primary adaptive routes for undesirable traits.

## INTRODUCTION

Experimental evolution is an essential tool for investigating adaptive walks, clonal dynamics, competition and fitness, and the genetic underpinnings of complex traits. One question experimental evolution enables us to explore is how often given the same conditions and selective pressures organisms will follow the same adaptive route (Gould’s “tape of life”) (Gould 1990; Orgogozo 2015). A primary platform for performing evolution experiments in the laboratory is the chemostat, a continuous culture device invented in 1950 by Monod (Monod 1950) and by Novick and Szilard (Novick and Szilard 1950). In a chemostat, new media is added and diluted at the same rate, maintaining constant growth conditions. Chemostat experiments have provided insight into the mechanisms of genome evolution and adaptation to a variety of selection pressures (reviewed in Gresham and Hong 2015). However, chemostats have been limited in their utility due in part to frequent selection for biofilms and cell aggregation, which have been observed since the advent of the chemostat and are thought to evolve due to selection by the physical constraints of the culture vessels. In 1964, Munson and Bridges recorded a selective advantage in a bacterial subpopulation that adhered to the wall of a continuous culture device (Munson and Bridges 1964). Topiwala and Hamer followed up on these findings in 1971 and suggested that encouraging this phenotype could actually lead to increased biomass output (Topiwala and Hamer 1971), an idea that has enjoyed success in subsequent years: chemostats are such a successful system for growing biofilms that they are often used to grow biofilms intentionally by supplying additional substrates to encourage biofilm development (Poltak and Cooper 2011).

In the context of experiments concerning traits unrelated to wall growth and aggregation, however, the ease of biofilm evolution in chemostats represents a significant problem. Since wall growth and aggregation phenotypes develop as an adaptation to the experimental vessel itself, they develop regardless of the intended selective pressures in any given experiment. The evolution of wall growth and cell aggregation inside the continuous culture vessel seeds competing subpopulations, differentially restricting nutrient access for aggregating cells and skewing the likelihood of dilution (Smukalla et al. 2008; Fekih-Salem et al. 2013), both variables that should be fixed. Thus, developing a strain with reduced potential for evolving biofilm-related traits in this type of experimental system has many practical benefits.

Combating biofilm-related traits is also important in industry and medicine. Biofilm formation is a cell-surface adhesion trait that enables pathogenic organisms to persist on the surfaces of medical devices and even colonize human tissues (Kojic and Darouiche 2004; Verstrepen, Reynolds, and Fink 2004). Flocculation, a related cell-cell adhesion phenotype (Guo et al. 2000), is a mechanism by which yeast can survive stresses including treatment from antimicrobial compounds (Smukalla et al. 2008; Stratford 1992), with the cells on the interior of the floc physically protected from chemical treatments that more easily kill the outer layer of cells. This makes flocculation a problematic trait from a health perspective, and illustrates the importance of better characterizing the genetic basis of complex biofilm-related traits.

Cell aggregation, which we define here as an umbrella term to include both flocculation and mother/daughter separation defects (Stratford 1992), has dozens of known contributing genes identified by QTL mapping, deletion collection, genetic screen, and linkage analysis studies (Lee, Magwene, and Brem 2011; Granek et al. 2013; Brem 2002; Ryan et al. 2012; Taylor and Ehrenreich 2014; H. Y. Kim et al. 2014; Roop and Brem 2013; Borneman et al. 2006; Ratcliff et al. 2015; Palecek, Parikh, and Kron 2000; Verstrepen et al. 2005; Cullen 2015; Taylor et al. 2016; Brückner and Mösch 2012). Given the extensive list of genes involved in aggregation that could potentially contribute to its evolution in a chemostat, our primary interests in this study were determining across many evolution experiments whether the genes involved in the evolution of aggregation were expected or novel, and ascertaining whether aggregating clones all achieved this final phenotype through one primary or many equally favored adaptive routes.

To ask how yeast evolve aggregation, we used multiplexed parallel evolution experiments coupled with genetics and whole genome sequencing to determine the causal mutations in 23 aggregating clones isolated from evolution experiments that ran 300 or more generations. Despite the known genetic complexity of aggregation, most of the causal mutations appeared to operate through a favored adaptive route: activating flocculation gene *FLO1.* Blocking this favored route by deleting *FLO1* significantly reduced incidence of flocculation in further evolution experiments, demonstrating the efficacy and potential of data-driven strain engineering, even for complex traits.

## MATERIALS AND METHODS

### Strains and media used in this study

The ancestral strain for all evolved strains used in this study was *Saccharomyces cerevisiae* laboratory haploid *MAT***a** strain FY4 (S288C), and backcrossing experiments were conducted using its isogenic *MATα* counterpart FY5. Standard growth medium for overnight liquid cultures and agar plates used in this study was yeast extract peptone dextrose (YEPD) media, with 2% glucose and 1.7% agar for plates. Glucose-limited, sulfate-limited, and phosphate-limited liquid media and plates were prepared as in Gresham *et al* (Gresham et al. 2008) and detailed media recipes are available at http://dunham.gs.washington.edu/protocols.shtml.

To construct a *flo1* knockout strain, KanMX was amplified from the *FLO1* locus in the *flo1* strain from the yeast knockout collection (Giaever et al. 2002) using primers CJA009F/R (Table S4). The PCR reaction was cleaned using a Zymo DNA Clean and Concentrator kit and DNA concentration was quantified with a Qubit fluorometer. Strain FY4 (S288C) was transformed with 1ug of the amplicon in 75µl of 1-step buffer (50% PEG4000 (40% final), 2M LioAc (0.2M final), 1M DTT (100nM final), salmon sperm carrier DNA) at 42C, and transformants were selected for G418 resistance. The *flo1::KanMX* strain was verified using Sanger sequencing.

### Multiplexed chemostat evolution experiments

The first set of evolved clones was generated from 96 evolution experiments, conducted with laboratory strain FY4. The experiments were split equally between three nutrient limited conditions, 32 each of glucose, sulfate, and phosphate limitation, and organized into six blocks of 16 vessels maintained at 30°C. The evolution experiments were set up and media was prepared according to (Miller et al. 2013), with minor modifications. Sampling was conducted daily. The dilution rate was maintained in a target pump setting range of 0.16 and 0.18 volumes/hour, and generations elapsed were calculated as (1.44)*(time elapsed)*(dilution rate). Total generations were calculated as the cumulative sum of these individual times. One vessel was lost to pinched pump tubing that obstructed its media supply, for a final number of 95 evolution experiments. The remaining 95 evolution experiments were terminated at ~300 generations. Throughout the experiment, vessels were monitored for evidence of wall sticking and aggregation, and in this initial experiment both traits were scored together. In later experiments, we scored these traits separately. 12/32 phosphate-limited, 18/32 glucose-limited, and 3/31 sulfate-limited populations demonstrated evidence of aggregation or wall sticking, and we selected 9 phosphate, 11 glucose, and 3 sulfate-limited populations for further analysis.

The comparison between *flo*1 knockout and wild-type strains was conducted using 64 glucose-limited chemostats run as above. Within each 16-vessel block, wild-type strains and knockout cultures were set up in alternating rows of 4. Up to 150 generations, sampling was conducted once weekly. Cultures were monitored daily for evidence of contamination, flocculation, and colonization in any of the media or effluent lines. After 150 generations, samples were stored twice weekly, and microscopy images for all cultures were saved once weekly. At the final timepoint, microscopy images were collected on all cultures. Clumps from the bottom of the culture or rings adhering to the vessel walls were collected with long sterile cotton swabs and resuspended in media and glycerol for storage. The final populations were plated on YPD to check for contamination and replica-plated onto G418 to validate the presence of the *KanMX* marker in only the expected *flo*1 knockout populations.

### Clone isolation

Colonies were struck out from glycerol stocks of the final time point of each experiment, inoculated into liquid culture and grown overnight at 30°C. From overnight cultures that displayed a clumping and/or settling phenotype, single cells were isolated using micromanipulation on a Nikon Eclipse 50i dissecting microscope, allowed to grow into colonies, screened for the phenotype in an overnight liquid culture of the appropriate nutrient-limited media, and saved at -80°C in glycerol stocks.

### Whole Genome Sequence analysis

Genomic DNA for each clone was extracted using a Zymo YeaStar genomic DNA kit, checked for quality using a NanoDrop ND-1000 spectrophotometer, and quantified using an Invitrogen Qubit Fluorometer. Genomic DNA libraries were prepared for Illumina sequencing using the Nextera sample preparation kit (Illumina) and sequenced using 150bp paired-end reads on an Illumina HiSeq. Ancestral DNA was prepared using a modified Hoffman-Winston preparation (Hoffman and Winston 1987).

Average sequence coverage from WGS of the clones was 97x. The reads were aligned against the genome sequence of sacCer3 using Burrows-Wheeler Aligner version 0.7.3 (Li and Durbin 2009). PCR duplicates were marked using Samblaster version 0.1.22 (Faust and Hall 2014) and indels were realigned using GATK version 3.5 (McKenna et al. 2010). For SNV and small indel analysis, variants were called using the bcftools call command (Li and Durbin 2009). SNVs/indels were filtered for quality and read depth, and mutations unique to the evolved clones were identified, annotated with a custom Python script (Pashkova et al. 2013), and verified by visual examination with the Integrative Genomics Viewer (IGV) (Robinson et al. 2011). This analysis revealed an average of three high quality SNVs/indels per clone after filtering, with a maximum of 16. Complete sequencing data for all of these clones is available under NCBI BioProject PRJNA339148, BioSample accessions SAMN05729740-5729793. Structural variants were called using lumpy (version accessed on 20160706) (Layer et al. 2014), and copy number variants were called using DNAcopy (Seshan and Olshen 2015) on 1000bp windows of coverage across the genome.

The deletion in gene *MIT1* was validated in clones YMD2694 and YMD3102 using PCR (primers EH053PF/PR) (Table S4) and Sanger sequencing. Validation of other mutations is described below.

### Microscopy and validation of separation defects

Strains were grown overnight in 5mL YEPD liquid culture at 30°C. 5µl of culture was examined microscopically at 150X magnification and photographed using a Canon Powershot SD1200 IS digital camera. Images were scored for evidence of mother-daughter separation defects, which were identified in two of the clones, YMD2680 and YMD2689. To validate the separation defects, calcofluor white was added to 1x10^7^ cells at a concentration of 100µg/mL, pipetted to mix, and incubated in the dark for 5 minutes or more. Cells were pelleted at 13200 rpm for 1 minute and the supernatant was removed. The pellet was then washed vigorously with 500µl water three times and re-suspended in 50µl water. Bud scars were visualized using a DAPI filter at 630X magnification.

To validate true flocculation in the remaining clones, the evolved clones and ancestral strain were inoculated into 100µl YEPD cultures in two replicates in a round-bottom 96-well plate and grown overnight at 30°C without agitation. Cultures were resuspended by pipetting and 5µl of culture was examined microscopically at 150X magnification and imaged. Cells were pelleted and the supernatant removed by pipetting, and one replicate was re-suspended in 100µl water and the other in 100µl 4mM EDTA. Each replicate was pipetted ten times and then examined microscopically and imaged. After ~50 minutes, replicates were re-suspended again by pipetting five times, and examined microscopically and imaged again.

### Quantitative settling assay

Settling analysis was conducted according to the protocol described in Hope and Dunham 2014 (Hope and Dunham 2014). Briefly, each evolved clone or backcross segregant was grown in 5mL YEPD for 20 hours at 30°C; strain YMD2691 and its segregants are slow growing so an additional replicate was completed for these segregants with 30 hours of growth. Each culture tube was vortexed and then placed over a black background to settle for 60 minutes, with photos taken of the settling culture at time zero immediately after vortexing and at time 60 after an hour of settling. Images were converted to black and white in Picasa version 3.9.141.306 and analyzed in ImageJ version 1.47v (Abramoff, Magalhães, and Ram 2004). The settling ratio (percent of tube cleared at 60 minutes) was calculated as in Hope and Dunham 2014 (Hope and Dunham 2014). Three replicate measurements were taken on each image of the evolved clones, and a single measurement was made for the segregants.

### Backcrossing and settling segregation patterns

All clones except YMD2680 and YMD2689, which had separation defects, were backcrossed to strain FY5. An average of 16 full tetrads per cross were dissected, with additional dissections for crosses with clones YMD2678 (24 tetrads total) and YMD2697 (38 tetrads total). Segregants were inoculated into 100µl YEPD in round-bottom 96-well plates and grown overnight at 30°C without agitation. Plates were re-suspended by gentle pipetting and allowed to settle without agitation for 15 minutes, when they were photographed and scored for settling ability.

### Bulk Segregant Analysis

Crosses for clones YMD2684, YMD2686, YMD2687, YMD2696, YMD2697, YMD2698, and YMD2699 were not utilized for BSA after this point as it was determined that they harbored the Ty insertion in the *FLO1* promoter; four strains that harbored this insertion (YMD2681, YMD2683, YMD2685, and YMD2690) were included in BSA to verify causality for the Ty insertion. The cross with clone YMD2701 was also not included for BSA because it did not segregate the settling trait 2:2. The nine remaining strains without a *FLO1* promoter Ty element insertion or separation defect were analyzed using BSA. Segregants were binned into two pools of cells according to phenotype (settling or non-settling). Cells were pelleted, washed once with 500µl water, transferred to a 2mL lock-top eppendorf tube, pelleted again, decanted, and frozen at -20°C until DNA extraction. Genomic DNA was extracted using a modified Hoffman-Winston preparation (Hoffman and Winston 1987). Sequencing libraries were prepared using Nextera library preparation protocols as described for the original clones. To identify causal mutations, BSA pools were analyzed similarly to the evolved clones but filtered individually by sample. For each sample, mutations present in both the settling and non-settling pool were removed. Mutations present at an allele frequency of 1 were determined to be causal.

### Identification of Ty element insertion location and element type

A Ty insertion in the promoter of *FLO1* was identified in 12 of the evolved flocculent clones by visual examination in IGV and split read analysis tool retroSeq (Keane, Wong, and Adams 2013). These insertions were verified as full-length using PCR with primers CJA007F/R (Table S4). In some cases, an exact breakpoint was determined using the program lumpy, but for other samples the Ty element insertion location was determined by visual examination in IGV. All insertions placed the Ty element in reverse orientation with respect to the *FLO1* gene, determined by manual analysis of the mapping orientation of split reads.

A 2.1kb region upstream of *FLO1* and including the start of the ORF was amplified using PCR with Phusion polymerase and primer pair CJA007F/R (Table S4) for each clone with a Ty insertion identified in WGS. The presence of a Ty element insertion leading to a 6kb expansion was verified on a 1% agarose gel. PCR verification of the insertion failed in three clones, YMD2681, YMD2683, and YMD2697. The PCR reactions were cleaned using a Zymo DNA Clean and Concentrator kit and eluted in 100µl of water. Ty1 contains two EcoRI sites not shared with Ty2, and Ty2 contains a unique BamHI site missing from Ty1; these features facilitate classification of Ty type by restriction digest. The cleaned amplicons were split into two restriction enzyme digest reactions, one with EcoRI and the other with BamHI (New England BioLabs). A model of the amplified region was created in sequence analysis software Ape (“ApE - A Plasmid Editor” 2016), with a Ty insertion in the middle of each hot spot insertion region: Ty insertions were observed between 95 and 156bp and between 394 and 470bp upstream of the *FLO1* ORF, so for the close insertion model a Ty1 was added at 125bp from the ORF and for the far insertion model at 432bp from the ORF. Predicted cutting with EcoRI for the close insertion site yielded four bands at 208, 1408, 2344, and 4118bp, and for the far insertion site four bands at 208, 1408, 2651, and 3811bp. We observed the three longest bands as predicted on a 1% agarose gel following the restriction digests, with distinct size differences between the mid and high bands for clones with known close and far insertions; for all evolved clones successfully analyzed, the insertion was classified as a Ty1. Predicted banding patterns for cleavage with BamHI in the region were also consistent with Ty1 elements. As a positive control, a known Ty2 element was amplified from the S288C genome using primers EH054PF/PR (Table S4) and the banding patterns that would be present for a Ty2 element with BamHI and EcoRI digests were confirmed.

### Crosses to determine *FLO1* dependence of mutations in *ROX3, CSE2*, and *MIT1*

*MATα* segregants of clones with mutations in *CSE2* (YMD2678), *ROX3* (YMD2691), and *MIT1* (YMD2694) were crossed to a *flo1* knockout strain to facilitate examination of the phenotype of the double mutant progeny, recorded based on the settling ratio of segregants in 16-18 tetrads per cross. Mating types of segregants were verified using a standard halo mating assay (protocol available at http://dunham.gs.washington.edu/protocols.shtml).

### Additional analyses for secondary modifiers in clones YMD2683 and YMD2690

Two strong candidates for clones with secondary modifiers were YMD2683, with an elongated cell morphology, and YMD2690, with an expansion of the internal repeats in *FLO11.* All of the segregants screened in the quantitative settling assay for clone YMD2683 were also tested for mutations in genes *HSL7, IRA1, VTS1*, and *TCP1* using primers EH045PF-EH048PR (Table S4) and sent for Sanger sequencing by Genewiz. Microscopy was performed on all of the segregants from the YMD2683 cross and nine additional segregants were selected based on cell morphology (two with round suspended cells, two with long suspended cells, two with round flocculent cells, and three with long flocculent cells); all were analyzed with the quantitative settling assay and sequenced for mutations in *HSL7* and *IRA1.*

For seven settling segregants from the backcross with clone YMD2690, the *FLO11* internal repeat region was amplified using primers EH030PF/PR (Table S4) and results were examined on a 1% agarose gel. For the same segregants, the region of *HOG1* surrounding a premature stop in the clone was amplified using primers EH052PF/PR (Table S4) and sent for Sanger sequencing.

## RESULTS

Experimental evolution studies using continuous culture systems have suffered from small sample sizes in the past, a challenge that has been addressed through our multiplexed miniature chemostat system (Miller et al. 2013). In a previous study designed to test changes in fitness in response to different nutrient limitations, we ran 96 miniature chemostats under three different nutrient limitations for 300 generations (Miller, A.W., in preparation). We observed that by 300 generations 34.7% had gained a visible cell aggregation phenotype not present in the ancestral strain, an S288C derivative that cannot aggregate due to a nonsense mutation in transcription factor Flo8 (Liu, Styles, and Fink 1996).

### Majority of aggregating clones demonstrate characteristics of true flocculation

We selected clones for further study from the 23 populations with a strong aggregation phenotype. We conducted a number of phenotypic and genotypic analyses on the selected clones in order to determine how each strain had independently evolved the ability to aggregate. We quantified the settling ability of the evolved clones compared to the ancestral strain (Fig. 1) (Table S1), a metric that describes the primary phenotype of interest in these experiments. We also examined the cellular morphology of all evolved clones microscopically and determined that two of the 23 clones (YMD2680 and YMD2689) show a cellular chaining phenotype indicative of a mother-daughter separation defect, while the remaining clones had aggregating round cells characteristic of cell-cell adhesion and true flocculation (Fig. S1). We confirmed the bud separation defect in YMD2680 and YMD2689 using calcofluor white staining (Fig. S2), which preferentially stains the increased chitin present at yeast bud scars (Pringle 1991). To further distinguish separation defects from flocculation, we treated the evolved clones with a de-flocculation buffer containing a chelating agent, EDTA; true flocculation is facilitated by calcium ions and reversible, while separation defects are not (Stratford 1989; Liu, Styles, and Fink 1996). We verified that all clones excluding YMD2680 and YMD2689 exhibit true flocculation that is reversible upon treatment with EDTA (Fig. S3).

**Figure 1:**
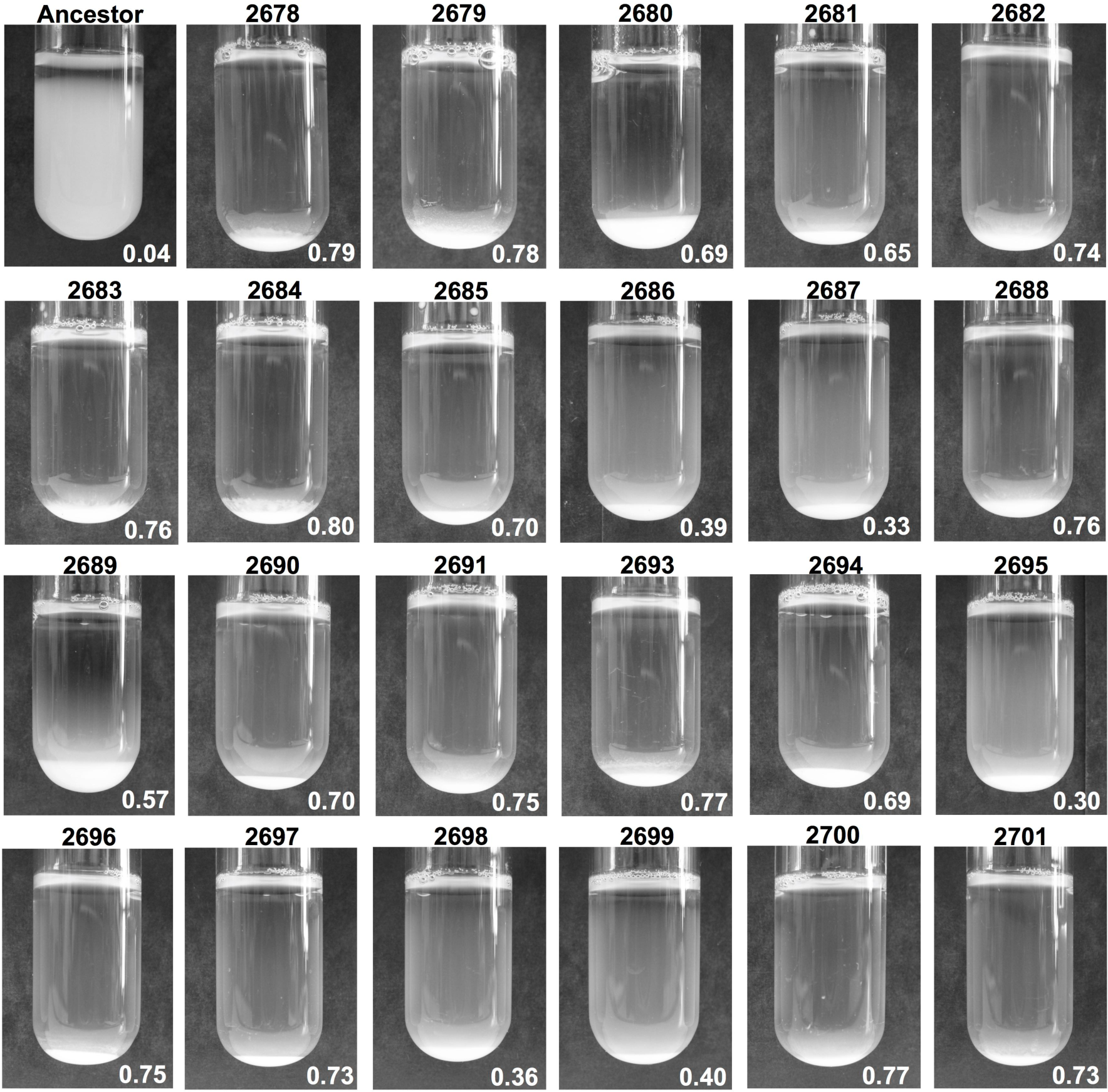
Quantitative settling of aggregating evolved clones. Images of the 60-minute settling time point for ancestral strain FY4 and 23 evolved clones with aggregation trait. Cultures were grown to saturation in 5mL YEPD liquid media. Settling ratio values are shown in bottom right of each image; ratios are the mean of three measurement replicates on the image shown.

### Mutations in *FLO1* promoter and genes *TUP1* and *ACE2* are primary adaptive routes to aggregation

We performed Whole Genome Sequencing (WGS) on the 23 clones from generation 300 of the evolution experiments and analyzed the resulting sequence data to identify Single Nucleotide Variants (SNVs), small insertions or deletions (indels), Copy Number Variants (CNVs), and structural variants (Table S2, Materials and Methods). We developed a list of candidate genes likely to contribute to the evolution of aggregation phenotypes (Table S3) from 17 different papers examining biofilm and cell aggregation related-traits, and several of the SNVs identified in our clones were in candidate genes (e.g. *ACE2, HOG1, TUP1).* We did not identify any instances of reversion of the ancestral point mutation in transcription factor gene *FLO8.*

In both clones harboring separation defects, we discovered short insertions and deletions in the transcription factor gene *ACE2*, both of which cause a shift in the reading frame and introduction of a premature stop codon. These results are consistent with prior literature showing that loss of function mutations in this gene cause settling/clumping phenotypes in other experimental evolution scenarios (Ratcliff et al. 2015; Voth et al. 2005; Oud et al. 2013; Koschwanez, Foster, and Murray 2013). Furthermore, *ace2* null mutants have the characteristic cell separation defect that we observed in our clones. We consider this *ACE2* mutation causative of the aggregation phenotype in these two clones (Table 1), with possible modification by *BEM2*, a gene involved in bud emergence that is also mutated in both clones (Bender and Pringle 1991; Y.-J. Kim et al. 1994).

**Table 1:** Causal mutations for the aggregation phenotype in 23 evolved clones.

| CLONE<br>(YMD) | NUTRIENT<br>LIMITATION | CAUSAL<br>GENE | SYSTEMATIC<br>NAME | MUTATION TYPE |
| --- | --- | --- | --- | --- |
| 2678 | G | *CSE2* | YNR010W | Ty insertion in ORF at R137 |
| 2679 | S | *TUP1* | YCR084C | Stop-gained Q181* |
| 2680 | G | *ACE2* | YLR131C | S115 indel (2bp deletion);<br>premature stop introduced |
| 2681 | G | *FLO1* | YAR050W | Ty in promoter (156bp upstream of<br>ORF) |
| 2682 | P | *TUP1* | YCR084C | Q107-P143 deletion in ORF<br>(106bp); premature stop introduced |
| 2683 | G | *FLO1* | YAR050W | Ty in promoter (139bp) |
| 2684 | S | *FLO1* | YAR050W | Ty in promoter (127bp) |
| 2685 | G | *FLO1* | YAR050W | Ty in promoter (397bp) |
| 2686 | P | *FLO1* | YAR050W | Ty in promoter (151bp) |
| 2687 | P | *FLO1* | YAR050W | Ty in promoter (156bp) |
| 2688 | P | *TUP1* | YCR084C | Ty insertion in ORF at L341 |
| 2689 | G | *ACE2* | YLR131C | L192indel (1bp insertion);<br>premature stop introduced |
| 2690 | G | *FLO1* | YAR050W | Ty in promoter (449bp) |
| 2691 | G | *ROX3* | YBL093C | Stop-gained C138* |
| 2693 | G | *TUP1* | YCR084C | Stop-gained Q101* |
| 2694 | S | *MIT1* | YEL007W | L552-M585 deletion in ORF<br>(101bp); premature stop introduced |
| 2695 | P | *FLO9* | YAL063C | 6.2kb deletion in promoter (229bp<br>upstream of ORF) |
| 2696 | G | *FLO1* | YAR050W | Ty in promoter (470bp) |
| 2697 | P | *FLO1* | YAR050W | Ty in promoter (95bp) |
| 2698 | G | *FLO1* | YAR050W | Ty in promoter (394bp) |
| 2699 | P | *FLO1* | YAR050W | Ty in promoter (102bp) |
| 2700 | P | *TUP1* | YCR084C | V592 indel (27bp deletion); in frame |
| 2701 | P | *FLO1* | YAR050W | Ty in promoter (152bp) |

The gene in which or in front of which the causal mutation was found is identified here, along with the type of mutation we recorded. Also shown is the nutrient limitation in which the clones were evolved: G, S, or P for glucose-limited, sulfate-limited, and phosphate-limited, respectively. Positional information about SNVs and indels is exact; other values shown are approximate (Materials and Methods).

In the 21 flocculent clones, the most common mutation we identified was a full-length insertion of a yeast transposable (Ty) element in the promoter region of *FLO1.* We saw this insertion in 12 of our clones, distributed in two hotspot regions between 95 and 156 bp and 394 and 470 bp upstream of the *FLO1* start codon (Fig. 2, with regulatory information from (Fichtner, Schulze, and Braus 2007; Fleming et al. 2014; Basehoar, Zanton, and Pugh 2004)). Sequence analysis narrowed the type of Ty element in these insertions to Ty1 or Ty2, and diagnostic PCR and restriction digestion of nine inserts confirmed they were all Ty1 elements. In *FLO1* overexpression, localization, and deletion studies, *FLO1* has been shown to cause flocculation (Guo et al. 2000; Bony, Barre, and Blondin 1998; Smukalla et al. 2008); notably, Smukalla et al demonstrated that GAL-induced expression of *FLO1* in S288C, the background strain for these evolution experiments, induces flocculation, which supports the role we observe for *FLO1* regulation.

**Figure 2:**
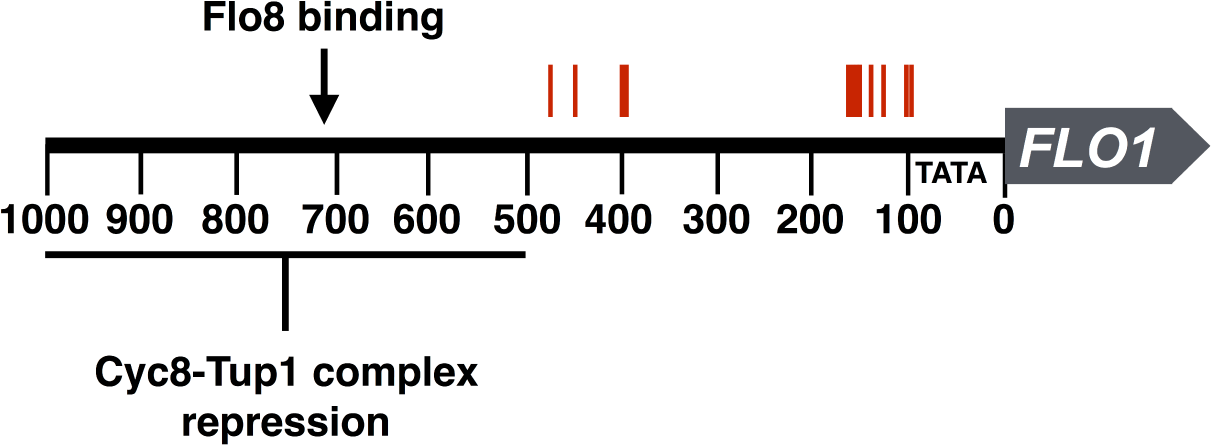
Ty element insertion sites cluster in two regions of the *FLO1* promoter. The region 1kb upstream of *FLO1* is shown with the insertion site positions of Ty elements observed in 12 evolved clones in red. Locations shown in this figure serve to demonstrate the primary regions of insertion only; for best estimates of exact insertion locations see Table 1. Flo8 binding site and Tup1-Cyc8 repression information adapted from (Fichtner, Schulze, and Braus 2007; Fleming et al. 2014). The TATA box is shown at 96 bp from the start of the open reading frame, as in (Basehoar, Zanton, and Pugh 2004).

In the remaining nine clones, we identified several SNVs and larger insertions and deletions in candidate genes, including *TUP1, FLO9, IRA1*, and *HOG1*, and many more in non-candidate genes (Table S4). Five clones harbored likely loss-of-function mutations in candidate *TUP1:* two stop-gained SNVs in clones YMD2679 and YMD2693; one 27 bp deletion in YMD2700; one 100 bp deletion in YMD2682; and one Ty element insertion in YMD2688. *TUP1* is a general repressor (Carrico and Zitomer 1998; Z. Zhang, Varanasi, and Trumbly 2002), but also a repressor of *FLO1* (Fleming et al. 2014), and loss-of-function mutations in this gene have been associated with flocculation since 1980 (Williams and Trumbly 1990; Teunissen, van den Berg, and Steensma 1995; Lipke and Hull-Pillsbury 1984; Stark, Fugit, and Mowshowitz 1980). The frequently observed mutations in *TUP1* could function to de-repress *FLO1* or any number of other candidate genes. In a different clone, YMD2695, we identified a 6.2 kb deletion from 229 bp to 6.4 kb upstream of flocculin gene *FLO9.* We also identified high confidence mutations not previously associated with aggregation in nearly all clones.

### Bulk Segregant Analysis verifies causal mutations in novel genes

Because of the number of high confidence mutations in each clone, we could make hypotheses about causality. To test causality and examine the genetic complexity of the trait in each clone, we turned to a different method, bulk segregant analysis (BSA). We backcrossed the 21 evolved flocculent clones to a non-flocculent strain isogenic to the ancestor but of the opposite mating type. We excluded the two clones with separation defects because their causality was clear and their budding defect interfered with tetrad dissection. BSA leverages meiotic recombination and independent assortment to link a trait to a causal allele, which will be observed in all progeny with the phenotype of interest. In turn, unlinked non-causal alleles should assort equally between progeny with and without the phenotype (Brauer et al. 2006; Birkeland et al. 2010; Segrè, Murray, and Leu 2006) (Fig. 3A). Backcrossing also allowed us to estimate the genetic complexity of the trait: if two of four meiotic progeny have the phenotype and two do not, this indicates a single causal allele for the phenotype. We observed this 2:2 segregation pattern in 20 of the 21 evolved clones, and pooled and sequenced the progeny with and without the trait to identify which of the initial candidate alleles was causal.

**Figure 3:**
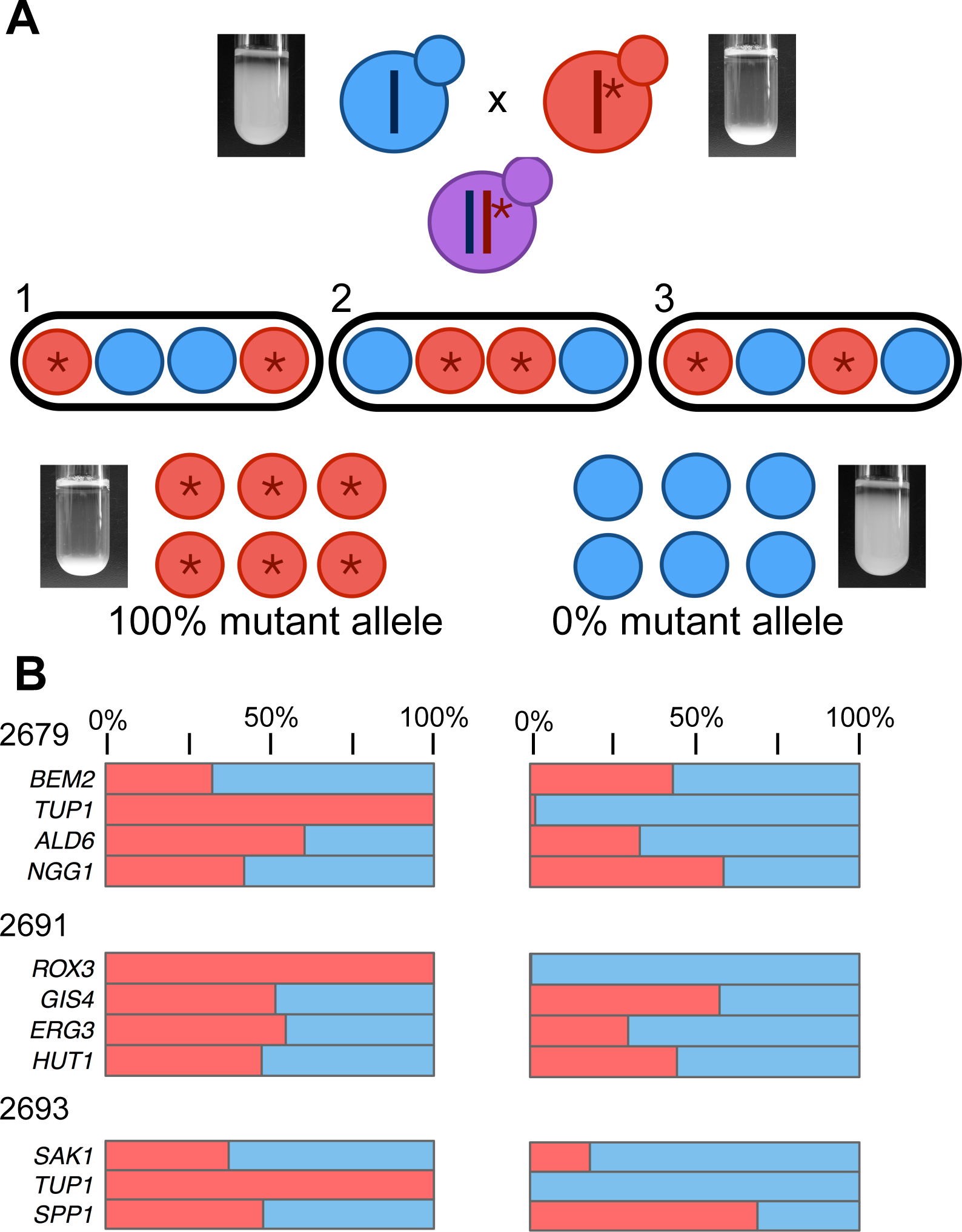
Bulk segregant analysis leverages recombination to identify mutations that co-segregate with the flocculation trait. **A)** An evolved clone with the phenotype of interest, shown here as settling in liquid culture, is backcrossed to the ancestral strain lacking the phenotype. Dissection of tetrads resulting from this cross reveals the segregation pattern of the trait among meiotic progeny, with 2:2 segregation (two segregants with the settling trait and two without) indicative of single gene control of the trait. Segregants with and without the trait are pooled and sequenced, and alleles that co-segregate with the trait are identified as causal. **B)** For three backcrosses, pooled sequencing results are shown for both pools of segregants, those with the settling trait on the left, and without the settling trait on the right. The strain identifier for the evolved clone in the cross is shown on the left, along with a list of candidate genes that had high quality Single Nucleotide Variant calls in the clone. The red bar shows the % of each of those candidate mutations seen in each pool, with mutations seen at 100% frequency identified as co-segregating with the trait and therefore causal.

We subjected all of our clones with genic mutations, including large insertions and deletions, to BSA analysis, and included four of the clones with a Ty element insertion in the *FLO1* promoter. The analysis pipeline (Materials and Methods) identified mutations at 100% frequency in the flocculent pools, and confirmed the causality of the *FLO1, TUP1, FLO9*, and *ROX3* mutations. BSA also confirmed the causality of mutations in *CSE2* and *MIT1*, genes not previously associated with flocculation (though both have been linked to related traits such as invasive growth and biofilm formation, see below). For the three evolved clones with causal SNVs, the frequency of each candidate in the flocculent and non-flocculent pools is shown in Fig. 3B and 3C; in each case, the causal mutant allele was at 100% frequency in the flocculent pool. Using the combined results of WGS and BSA, we were able to resolve the causal mutation for all 23 of the evolved clones, with a complete summary of our findings in Table 1.

### Functional *FLO1* is necessary for flocculation driven by *ROX3, CSE2*, and *MIT1* mutations

Given the large number of potentially activating mutations that we recovered in *FLO1*, we hypothesized that the causal *ROX3* and *CSE2* mutations we recorded also act through *FLO1*, via loss of repression. Several lines of evidence make *ROX3* a reasonable candidate repressor for *FLO1* and/or other *FLO* genes. Loss-of-function mutations in *ROX3* have been previously associated with flocculation (T. A. Brown, Evangelista, and Trumpower 1995) and also pseudohyphal growth, which is a trait related to haploid invasion and regulated by *FLO* genes (Guo et al. 2000). *ROX3* and *CSE2* both encode components of the RNA polymerase II mediator complex, which also includes Sin4, Srb8, and Ssn8, whose role in *FLO* gene repression is described in Fichtner *et al* (Fichtner, Schulze, and Braus 2007). Mutations in other components of Mediator have previously been shown to cause clumping (Koschwanez, Foster, and Murray 2013). In order to test the relationship between the *ROX3* and *CSE2* mutations and *FLO1*, we examined the ratio of settling to non-settling progeny in crosses between a *flo1* knockout strain and strains harboring the *CSE2* and *ROX3* causal mutations. 50% settling and 50% non-settling segregants compiled over all tetrads would indicate, for example, that both the *cse2 FLO1* and *cse2 flo1* segregants flocculate and that the function of the *cse2* mutation is not dependent on a functional *FLO1.* 25% settling and 75% non-settling segregants, and the presence of tetrads segregating 1:3 and 0:4, would indicate that the double mutant does not flocculate and a functional *FLO1* is required for the effect of the *cse2* (or *rox3*) mutation to be observed. We observed that the double mutants show a wild-type, non-flocculent settling phenotype, i.e., that *flo1* is epistatic to the other mutations. This indicates that *FLO1* is required for these mutations to have an effect and lends support to the hypothesis that Rox3 and Cse2 function as *FLO1* repressors in the wild-type strain.

Analysis of progeny with a *flo1* null mutation and the *MIT1* allele from YMD2694 revealed similarly that the *MIT1* mutation requires a functional *FLO1* to cause flocculation. *MIT1* is a known transcriptional regulator of flocculin genes *FLO1, FLO10*, and *FLO11*, and null mutants of *MIT1* exhibit reductions in hallmark biofilm-related traits including invasive and psuedohyphal growth and colony complexity (Cain et al. 2012), which are related to flocculation in S288C (Liu, Styles, and Fink 1996; Fichtner, Schulze, and Braus 2007). This role of *MIT1* in the literature suggests that the deletion we record is not a loss-of-function mutation, although it is not dominant. If the *MIT1* deletion in YMD2694 caused loss of function, we would expect to see a non-flocculent phenotype, as we confirmed is observed in a *mit1* deletion strain; instead, the deletion causes a flocculation phenotype, indicating that it serves in some way to enhance the function of *MIT1*. The deletion itself is out of frame and therefore results in a modified C-terminus of the protein, including a premature stop codon and truncation of the final product. From the extensive literature on the *MIT1* ortholog in *Candida albicans, WOR1*, we know that DNA binding activity is likely confined to the N-terminal portion of the protein, far from the mutation in this allele of*MIT1:* in *WOR1*, two DNA binding regions in the N-terminal portion of the protein are sufficient for full activity (Lohse et al. 2010; S. Zhang et al. 2014). *WOR1* and *MIT1* also both have a self-regulatory mechanism through a positive feedback loop, a potential mechanism for the enhanced function implicated by the mutation we observe (Cain et al. 2012; Zordan, Galgoczy, and Johnson 2006).

### Phenotypic variation suggests secondary modifiers influence flocculation

Though we identified the *FLO1* promoter Ty element insertion as the primary causal allele for the aggregation trait in 12 of our clones, we observed variation in the types of flocs produced in our preliminary microscopy of the clones (Fig. S1), and differences in settling even among all strains with a *FLO1* promoter insertion. These differences were not caused by Ty element direction, proximity, or type: all of the Ty elements we were able to validate with PCR and restriction site polymorphisms were of type Ty1. Secondary genetic modifiers of the flocculation trait are an alternative explanation for this phenotypic variation. To identify strains potentially carrying secondary modifier mutations, we examined the distribution of quantitative settling ratios across a subset of settling segregants for each cross (Fig. S4) (Table S1). Segregants without a modifier were expected to match the evolved parent settling phenotype, while a distribution of settling abilities would be seen as evidence of a potential modifier (Fig. S4).

One strong candidate for multiple alleles contributing to the aggregation phenotype was clone YMD2701, the only evolved clone that did not segregate the settling phenotype 2:2 during BSA. Sequencing analysis revealed this clone does have the *FLO1* promoter Ty1 insertion. We also identified an amplification of chromosome I in this clone, both copies of which have the promoter insertion, indicating that two copies of the causal allele are segregating in this backcross; this genotype is consistent with the segregation pattern we observed (Fig. S5). Within the segregant settling ratios, however, we did not observe this aneuploidy to be a modifier of the trait (Fig. S4).

In clone YMD2683, we identified a secondary modifier related to cell morphology. In our initial microscopy (Fig. S1), we observed that clone YMD2683 had an unusual elongated cell morphology, which we observed segregating in the backcross as well. Microscopy of segregants from this cross revealed four phenotypic classes: round, suspended cells; round, flocculent cells; long, suspended cells; and long, flocculent cells (Fig. 4A). Segregants from the backcross involving evolved clone YMD2683 had two different settling ratios, the weaker of which correlated with the round, flocculent morphology, while the stronger settling ratio correlated with the long, flocculent cell morphology (Fig. 4A, C). WGS of the original clone identified high quality SNVs in genes *IRA1, HSL7, VTS1*, and *TCP1.* PCR and Sanger sequencing of each of these genes in segregants from each phenotypic class revealed co-segregating missense mutations in *HSL7* and *IRA1* in all segregants with the long cell phenotype (examples in Fig. 4B), suggesting that one or both of these mutations is functioning as a secondary modifier to enhance the phenotype from the *FLO1* Ty element insertion. *HSL7* and *IRA1* are located only 13kb apart from each other on chromosome II, indicating that this co-segregation could be due to linkage rather than the contribution of both genes to the trait, though null mutations in each have been linked to abnormal cell morphology very similar to that of our strain (*HSL7*, (Kucharczyk et al. 1999; Fujita et al. 1999)) and flocculation *(IRA1*, (Verstrepen, Reynolds, and Fink 2004; Halme et al. 2004)).

**Figure 4:**
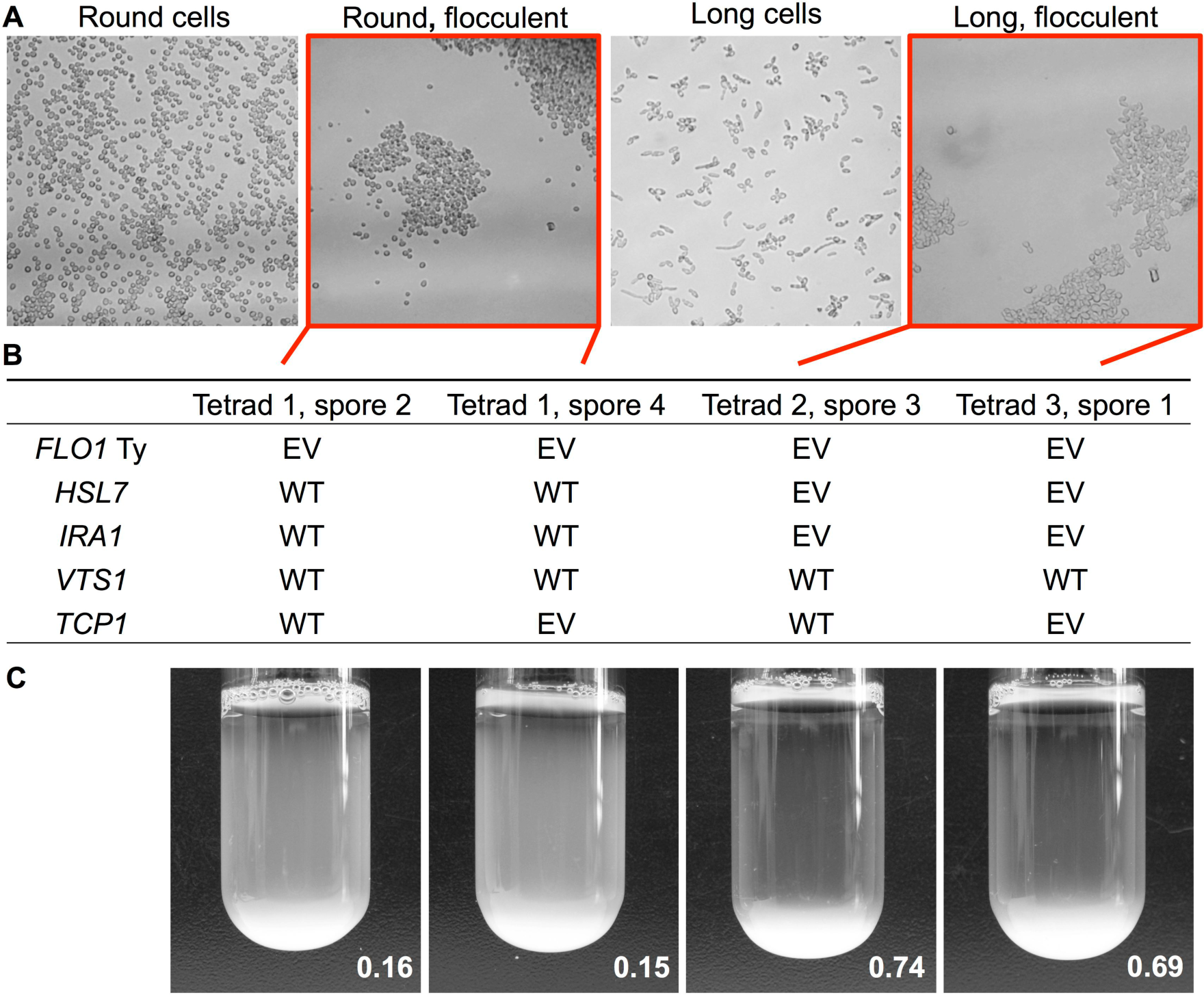
Evolved clone exhibits morphology-related secondary modifier of flocculation phenotype. **A)** Meiotic segregants show four phenotypic classes combining morphology and flocculation. Micrographs received additional processing (grey scale conversion, 20% increase in brightness, 20% increase in contrast) to better highlight the phenotypes. **B)** Sequencing results for four example flocculent segregants are shown, from two of the phenotypic classes in 4A: two segregants have round flocculent cells, and two with long flocculent cells. All segregants have the *FLO1* Ty insertion. The four candidate SNVs from the evolved clone were Sanger sequenced in the flocculent progeny and the match to the WT (S288C reference) or mutant/evolved base (EV) is recorded. **C)** Settling images and ratios for the four flocculent segregants that provided the sequencing data in 4B.

Another promising candidate for a secondary modifier was clone YMD2690: segregants from the backcross with this clone showed considerable variation in settling ratios, and the clone harbored a premature stop codon in candidate gene *HOG1* (Table S3), although a Ty element in the *FLO1* promoter was identified as the primary causal mutation. Using Sanger sequencing of settling segregants we determined that *HOG1* was not a secondary modifier of the trait. We conducted additional testing using primers from Zara *et al* (Zara et al. 2009) (Table S4) to target the repeat region in flocculin gene *FLO11* in clone YMD2690 and found evidence in this clone of a *FLO11* repeat expansion of approximately 1 kb in length. All flocculin genes have long arrays of internal tandem repeats (Verstrepen et al. 2005); expansions of the internal repeats in *FLO11* have been shown to cause phenotypic variability in biofilm-related traits, and natural isolates of yeast exhibit significant variation in the copy number of the repeats (Fidalgo, Barrales, and Jimenez 2008; Zara et al. 2009). However, this expansion also did not correlate with variations in strength of the segregant settling ratios, demonstrating that the presence of the *FLO11* expansion in addition to the Ty element insertion did not significantly affect the strength of the phenotype. PCR of all of the evolved clones revealed that only this clone had any evidence of repeat expansion in *FLO11.*

### Deleting *FLO1* increases time to evolve flocculation and reveals alternate adaptive routes

The results of our analyses of the evolved clones demonstrate a clear role for *FLO1* in the evolution of flocculation; not only do we see changes in the *FLO1* promoter, many of the other mutations we recorded are in genes encoding proteins that function to regulate *FLO1 (TUP1)* or participate in complexes that regulate *FLO1 (ROX3, CSE2).* We hypothesized that changes in the regulation of *FLO1* cause the flocculation phenotype in nearly all of the evolved clones, and that deleting *FLO1* would be a promising route for slowing the evolution of flocculation. Deleting a combination of *FLO* genes has been previously employed as a method to try to make lab strains easier to work with over long term experimental evolution (Voordeckers and Verstrepen 2015), and modification of the *FLO1* promoter has been effectively employed in biological circuits controlling flocculation (Ellis, Wang, and Collins 2009); however, it is unknown if specifically deleting *FLO1* would be effective on its own. We constructed a *flo1* strain and evolved 32 chemostat vessels of wild-type concurrently with 32 chemostat vessels of the *flo1* knockout strain, in glucose limited media for over 250 generations. Two knockout and one wild-type vessel were lost to contamination after generation 200. We monitored all vessels for evidence of aggregation and recorded eight wild-type and one knockout strain that developed aggregation during the course of the experiment, a statistically significant reduction *(p=0.01*, Fisher’s Exact Test). In order to determine the mechanism of the single aggregating *flo1* population, we performed WGS of a clone and found a Ty element insertion in the promoter of *FLO9.* In addition to the single knockout clone, we sequenced four wild-type strains that also evolved aggregation in the course of the experiment. Two of these harbored *FLO1* promoter Ty element insertions; another had a stop-gained mutation in *NCP1* that has not been verified as causal; and another had a deletion in *MIT1* exactly matching the deletion identified in the clone from the previous series of evolution experiments.

### *FLO1* deletion does not affect rate of evolution for unrelated traits

We expected that deleting *FLO1* would not impact the rate of evolution for unrelated traits, including wall sticking and separation defects, two other traits we monitored during the knockout evolution experiments. Cell-surface adhesion traits are more often associated with expression of *FLO11* (Guo et al. 2000; Verstrepen and Klis 2006), and we would not expect the frequency of evolving separation defects to be affected by changes to flocculation genes. For 32 of the vessels across both genotypes we recorded the occurrence of some amount of wall sticking two days before the final time point; eight of these we recorded as strong wall growth at the final evolution time point. The strong wall growth observations were split equally between WT and *flo1* knockout populations. Similarly, for mother-daughter separation defects, which we observed through microscopy of each of the final evolution time points, we recorded 12 strains with separation defects, five from the wild-type and six from the knockout evolution experiments, with an additional wild-type strain with an inconclusive microscopy phenotype (Fig. S6).

To explore the genetic origins of the wall sticking trait, we isolated clones with a strong wall sticking phenotype from six different populations, four from knockout experiments and two from wild-type experiments. Under the microscope, we observed that all six wall sticking clones harbored a separation defect. To determine if these were all caused by loss of function alleles of *ACE2*, we performed a complementation test using the *ace2* strain from the yeast deletion collection (Giaever et al. 2002). We determined that a loss-of-function mutation in *ACE2* was responsible for both the wall sticking and mother-daughter separation defects in five of the six clones. Despite this relationship, we did not observe a strong connection between wall sticking and separation defects on the population level, with 10 strains having only a separation defect, five having only strong wall growth, and only four populations having both phenotypes. The mechanism by which loss of function in *ACE2* facilitates wall sticking remains undetermined.

## DISCUSSION

Previous studies have successfully leveraged experimental evolution to understand the genetic contributors to complex traits (Voordeckers and Verstrepen 2015; Leu and Murray 2006; C. J. Brown, Todd, and Rosenzweig 1998; Hong and Gresham 2014). Evolution experiments have also contributed significantly to our understanding of how genomes evolve and the types of mutations typically observed in yeast grown in chemostats, including SNVs, CNVs, aneuploidy, and transposable element insertions (Dunham et al. 2002; Araya et al. 2010; Gresham et al. 2008; Adams, Julian 2004; Adams and Oeller 1986). In our study, we built on these concepts to identify the mutations contributing most to the evolution of cell aggregation, an industrially and medically relevant trait in addition to a practically useful one for facilitating laboratory work. We determined that in experimental evolution in continuous culture, loss-of-function mutations in *ACE2* are the most common contributors to the evolution of mother-daughter separation defects and wall growth, and mutations that change the regulation of *FLO1* are the most common evolutionary route to flocculation. The majority of causal mutations identified in this study occurred in candidate genes selected for involvement in aggregation traits based on previous literature, but two of the causal mutations were in genes not previously associated with flocculation *(CSE2, MIT1).* Both our identification of new genetic associations with flocculation and of one favored adaptive route to flocculation demonstrate the efficacy of using experimental evolution as a tool to better understand important complex traits.

This study also demonstrates the power of evolution experiments to determine which genes, among the many genes that are associated with complex traits like flocculation, most frequently contribute to adaptation under specific constraints. Despite the many possible candidates, we saw few of those genes identified in the evolved clones in this study. This finding is in keeping with other work in eukaryotes demonstrating favored adaptive responses, not just in the clear relationships between nutrient limitation and the amplification of nutrient transporters (C. J. Brown, Todd, and Rosenzweig 1998; Gresham et al. 2008) but also in response to stress treatments. In a more saturated screen we might start to see contributions from other candidate genes or pathways, but large screens have also revealed parallel adaptation. A study of 240 yeast strains under selective pressure from the antibiotic Nystatin revealed significant parallelism in mutational response through a single pathway (A.C. Gerstein, Lo, and Otto 2012) similar to the parallelism we discovered through *FLO1* regulation. The contributions of other candidate genes might also be revealed in a scenario in which an engineered strain has been modified to take away the primary adaptive routes we observed.

The primary mechanism of evolution we observed, a Ty element insertion in the *FLO1* promoter region, likely activates *FLO1* expression similarly to previous Ty element systems (Rothstein and Sherman 1980; Errede et al. 1984; Coney and Roeder 1988). The reverse orientation of the Ty element with respect to the open reading frame that we observed in all of our clones is the most common activating arrangement (Servant, Pennetier, and Lesage 2008; Boeke 2016), and the role of transposable elements in driving adaptive mutations has been well documented in yeast and other organisms (Chao et al. 1983; Tenaillon et al. 2016; Wilke and Adams 1992). Despite discovering one primary mechanism for evolving flocculation, we also show evidence for other genetic contributors modifying and enhancing the phenotype we observe. There is quantitative variation among settling segregants from crosses with our evolved clones (Fig. S4) particularly among strains with the *FLO1* promoter Ty element insertion, and we confirmed one example of a secondary modifier of the settling trait in clone YMD2683, in which a change in cell morphology enhanced the trait from the *FLO1* promoter Ty element insertion. Across other clones with trait variation there is potential to discover additional modifiers, both in the form of known candidate genes, including other *FLO* genes with internal tandem repeats, and in genes that have not previously been associated with flocculation.

Each causal mutation in our clones represents a new possible avenue for engineering to reduce aggregation. These could be simple changes, such as fusing genes like *ACE2* and *TUP1* that frequently acquire loss-of-function mutations to essential genes, or increasing their copy number or strain ploidy to increase the likelihood of “masking” deleterious recessive mutations (Otto and Goldstein 1992). They can also be iterative: deleting *FLO9* in the *flo1* background could even further reduce evolution of flocculation. Alternative strategies include reducing the mutation rate of these nondesirable mutations. The frequency at which we observe activating Ty elements driving flocculation also suggests future experiments aimed at reducing Ty element expression or mobility could be fruitful. Promising routes for reducing the Ty burden in evolution experiments include inhibiting Ty1 transposition (Xu and Boeke 1991) or utilizing different background strains. There is evidence that strain background contributes significantly to the likelihood of evolving flocculation in chemostat experiments. *Saccharomyces uvarum*, a budding yeast related to *S. cerevisiae* and often used in interspecific hybrid studies, has only Ty4 elements in its genome (Liti et al. 2005) and evolves flocculation more slowly than *S. cerevisiae* in chemostat experiments (Heil et al. 2016; Sanchez et al. 2016). Not only do different species of yeast have different Ty element burdens, natural isolates of *Saccharomyces cerevisiae* also provide strain-specific differences in Ty element burden (Bleykasten-Grosshans, Friedrich, and Schacherer 2013; Dunn et al. 2012) and a reservoir of variation in evolutionary potential which will be useful in future evolution experiments for studying flocculation and other complex traits.

Over the past six decades, experimental evolution in chemostats with yeast and bacteria has provided valuable insights into evolutionary dynamics and has proven to be a powerful tool for understanding complex traits. Now, with the advent of modern sequencing technology and common strain engineering methods, experimental evolution represents a promising direction for designing and testing strains with reduced (or increased) evolutionary potential. Evolution is gaining popularity as a tool for engineering: as just a few examples, in 2002, Yokobayashi *et al* used directed evolution to improve the function of a rationally designed circuit driving a fluorescent reporter (Yokobayashi, Weiss, and Arnold 2002), and evolutionary engineering is commonly used to improve carbon source utilization of industrial strains (Garcia Sanchez et al. 2010; Shen et al. 2012; Zhou et al. 2012). Evolution poses a challenge to strain engineering as well: loss, change, and breakage of engineered pathways confounds consistent usage (Renda, Hammerling, and Barrick 2014). Our study employs experimental evolution as a tool for engineering, but as a method both to design and to test new strains. We utilized evolution experiments as a means both to discover the genetic underpinnings of a complex trait with real-world applications, and to determine and eliminate the most successful adaptive route in order to generate a more amenable strain background for future experiments. This approach represents a promising engineering technique not just for flocculation and related traits but also for traits such as antimicrobial resistance that represent major challenges of our time.

## ACKNOWLEDGMENTS

We thank Noah Hanson, Monica Sanchez, Erica Alcantara, Michelle Hays, Bryony Lynch, Mei Huang, and Annie Young for experimental assistance. We also thank Aimée Dudley and Matthew Bryce Taylor for helpful comments on the manuscript, students participating in the Cold Spring Harbor Laboratories Yeast Genetics and Genomics Course in 2014, 2015, and 2016 for their contributions to the Bulk Segregant Analysis components of this project, and Maxwell W. Libbrecht for manuscript review and statistics consultation. This project was supported by NSF grant 1120425 and NIH grant P41GM103533. This material is based in part upon work supported by the National Science Foundation under Cooperative Agreement No. DBI-0939454. The CSHL Yeast Course is supported by NSF grant MCB-1437145. Any opinions, findings, and conclusions or recommendations expressed in this material are those of the author(s) and do not necessarily reflect the views of the National Science Foundation. CJA and CSH were supported by T32 HG00035. MJD is a Rita Allen Foundation Scholar and a Senior Fellow in the Genetic Networks program at the Canadian Institute for Advanced Research.

